# Identifying competition phenotypes in synthetic biochemical circuits

**DOI:** 10.1101/2022.03.22.485420

**Authors:** M. Ali Al-Radhawi, Domitilla Del Vecchio, Eduardo D. Sontag

## Abstract

Synthetic gene circuits require cellular resources, which are often limited. This leads to competition for resources by different genes, which alter a synthetic genetic circuit’s behavior. However, the manner in which competition impacts behavior depends on the identity of the “bottleneck” resource which might be difficult to discern from input-output data. In this paper, we aim at classifying the mathematical structures of resource competition in biochemical circuits. We find that some competition structures can be distinguished by their response to different competitors or resource levels. Specifically, we show that some response curves are always linear, convex, or concave. Furthermore, high levels of certain resources protect the behavior from low competition, while others do not. We also show that competition phenotypes respond differently to various interventions. Such differences can be used to eliminate candidate competition mechanisms when constructing models based on given data. On the other hand, we show that different networks can display mathematically equivalent competition phenotypes.

## I. Introduction

### A. Background

Living cells have the ability to perform sophisticated operations that include maintaining homeostasis against noise, responding appropriately to various input signals, constructing complex structures such as proteins, and adapting to novel environments. Reverse engineering the biochemical circuits responsible for implementing such operations has revealed various control mechanisms that include regulation of gene expression via transcription factors (TFs) and/or non-coding RNAs [1, 2]. This has inspired the development of engineering approaches that mimic natural circuits by inserting new synthetic circuits into cells to modify their behavior or create new functionalities. This has a wide range of applications that include immunotherapy, programmed micro-organisms for diagnostics and therapy, biofuel production, and many others [3–5]. Despite the great promise, numerous challenges exist. In particular, genetic circuits utilize common resources for transcription and translation such as RNA polymerases (RNAPs), ribosomes, tRNAs, and others. Insertion of new circuits increase the load on the cell’s reservoirs. This, in turn, can create indirect interactions that impede the proper functioning of the circuit, retard cellular growth, or lead to premature apoptosis [6, 7]. Several approaches have been proposed to ameliorate this problem, including dynamic control [8], orthogonal ribosomes [9, 10], forward engineering of the circuit to account for resource competition [11], and distributed computation [12, 13].

Several of the aforementioned approaches assume that it is possible to identify the mode of competition and the limited resources responsible for performance degradation. However, it is not always possible to infer the correct model of competition from the expression data of the circuit. Possible competition effects to account for include promoters competing for RNAPs, mRNAs competing for ribosomes, transcription factors competing for promoters, enzymes competing for substrates, substrates competing for enzymes, etc.

In this work, we identify *competition phenotypes,* i.e., features that could allow one to distinguish the scarce resource (or resources) responsible for performance deterioration. Are there *qualitatively different* types of competition effects? Are there *equivalent* effects that can be treated in a unified manner? Answers to these questions will help guide theoretical analysis as well the design of targeted interventions that mitigate undesirable effects through, for example, the use of feedback control to regulate the level of a scarce resource, or the optimization of appropriate circuit parameters. We will ask how different aspects of gene expression are impacted by two factors: (1) the level of the resource being shared –such as an activating or repressing transcription factor (TP), RNA polymerase (RNAP), or ribosomes– and (2) the level of competition from other biochemical species –such as other genes or mRNAs. We describe interventions that can be used to distinguish the different competition phenotypes. We also discover instances where, conversely, competition for different resources might result in mathematically equivalent competition prototypes.

### B. Problem Setup

#### Notation

Chemical species are denoted by non-italic large caps, time-dependent state variables are denoted by small caps, and constants are denoted by large cap italics. For example, a protein is denoted by Y, its time-dependent concentration is denoted by *y*(*t*) while its steady-state value is denoted by *Y*.

##### 1) The system

Consider a synthetic circuit with an external input *U* and internal state vector *x* which includes the concentrations of promoters, mRNAs, proteins, etc. The output is denoted by *y.* The circuit utilizes the free limited resource *r* which is also utilized by other *competing* (or interfering) circuits with inputs *I*_1_,…,*I_n_* and internal state vectors *z*_1_,…,*z_n_.* Figure 1 provides a pictorial representation of the system.

**Fig. 1.**
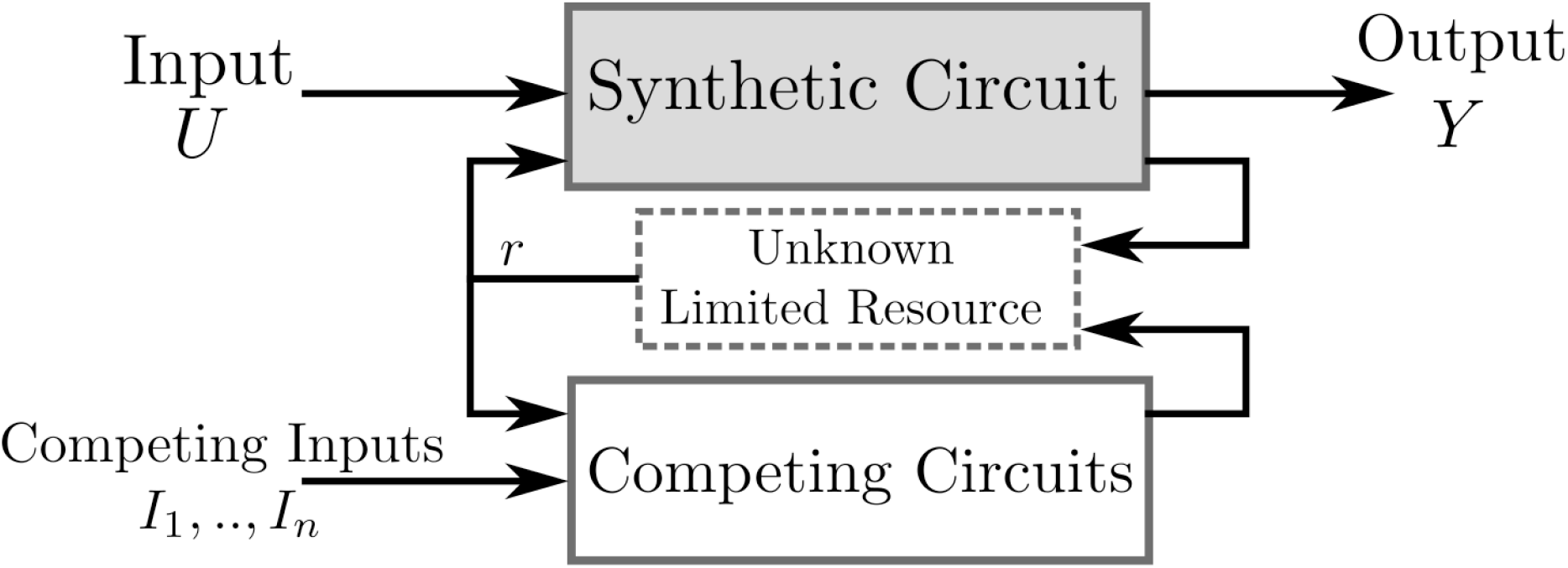
Framework for studying the problem of competition model discrimination.

We model the entire system by a system of ordinary differential equations structured as follows:

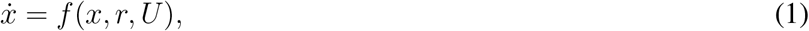

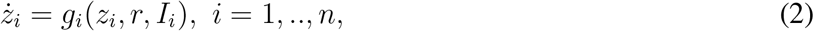

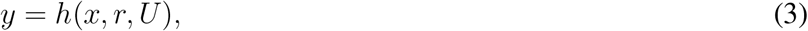

for some *C*^1^ vector fields *f, g*_1_,..,*g_n_*, and function *h*.

We assume that the limited resource is conserved. It partakes in the synthetic and competing circuits without being consumed or annihilated. Concretely, there exists nonnegative vectors *d*_0_,..*d_n_* of compatible dimensions such that

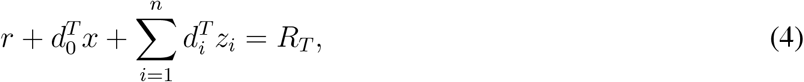

where *R_T_* is the total amount of the resource, and *r* is the free (available) resource. The time-evolution of the free resource is given by:

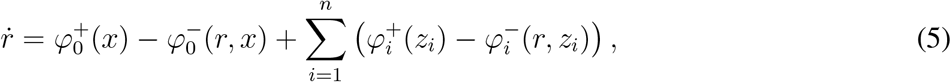

where 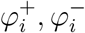, *i* = 0,.., *n* are functions which we call the inflow and outflow rates. In order for (1),(2),(5) to satisfy (4), the following relation is also satisfied: 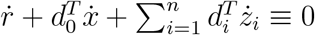.

In this paper, we perform our analysis at steady-state. We will assume that for each choice of the inputs *U, I*_1_,..,*I_n_* and total resource *R_T_*, there exists a steady-state (*X, Z*_1_,.., *Z_n_, R*) that is globally asymptotically stable.

After eliminating the intermediate variables, the corresponding steady-state output *Y* can be written as:

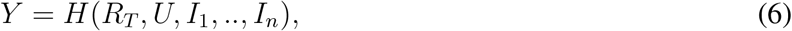

for some function *H*.

##### 2) Performance evaluation

The performance of a circuit with competition is compared to its performance with no competition. For this purpose, we introduce the *Competition-induced Performance Deterioration Ratio* (CPDR) *ρ* defined as:

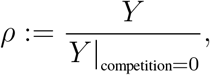

where the total competition is 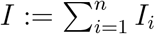.

##### 3) Problem Formulation

The formulation in (1)–(3) implicitly assumes that *r* is the limited resource, while the rest of the resources are abundant and they appear as kinetic constants in the functions *f, g*_1_,..,*g_n_, h*. Changing the identity of the limited resource will change the model (1)-(3). Our aim is to compare the qualitative differences in the steady-state input-output data that follow from the scarcity of different resources.

The paper is organized is follows. In section II, we discuss transcription/translation systems where the mRNA is protected by the ribosomes, while we drop that assumption in section III. Finally, we consider applications of the formalism to other bio-molecular contexts in section IV.

## II. Transcription/Translation model: Ribosome protects mRNA

Consider a synthetic genetic circuit in which a gene promoter is inserted in a cell, and it needs to be expressed in order to produce a desired output protein Y. The input to the circuit is the total concentration of the input promoter *U*, while the concentration of the unbound (free) promoter is denoted by *U_f_*.

We first assume that the mRNA molecules which are bound to the ribosome have a negligible rate of decay, which is a valid assumption in many situations [14]. The transcription and translation reactions can be then written as follows:

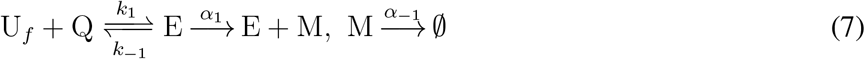

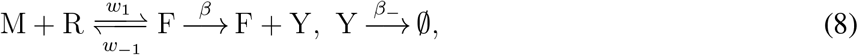

where Q denotes RNAP, R denotes the ribosome, E is the promoter-RNAP complex, and F is the mRNA-ribosome complex.

Free competing promoters I_*f,i*_, *i* = 1,.., *n* can bind to RNAP and produce mRNAs *M_i_* which can bind to the ribosome. This can be written as:

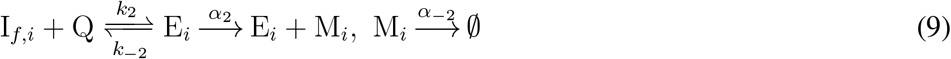

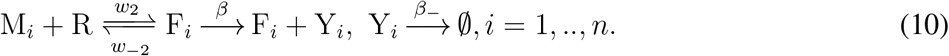

The promoters are conserved. Hence, we have:

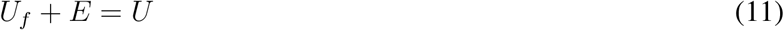

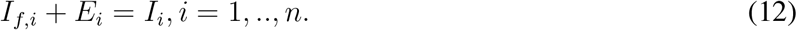

Let *K_j_* = *k_-j_/k_j_, A_j_* = *α_j_/α_–j_, W* = *w_–j_/w_j_, B* = *β/β_-_, j* = 1, 2 be RNAP Dissociation Ratio (DR), transcription ratio, ribosome DR, and protein expression ratio, respectively. The steady-states of the network (7)-(8),(9)-(10) can be computed by noting that the network is detailed-balanced, i.e. the forward and backward rates in each reaction are equal at steady state. Hence, the steady state values of the mRNAs *M, M_i_* are determined by the equilibrium values *A*_1_*E*, *A*_2_*E_i_, i* = 1,..,*n*, respectively, and are independent of the translation process. We get the following steady-state expressions:

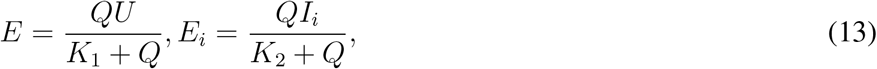

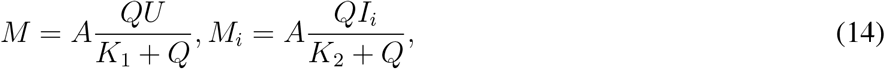

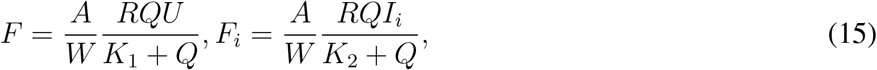

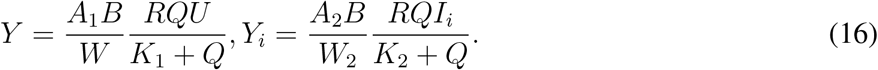

Note that *Q, R* are the levels of the *free* RNAP and ribosome, respectively.

In the formalism defined in §1.B and depending on the identity of the limited resource, Eq. (1) describes the dynamics of (7)-(8), while Eq. (2) describes the dynamics of (9). In order to write the output in the form (6), we will study several scenarios in which either the ribosome is limited, or RNAP is limited. We keep the other resource abundant to isolate the effects of the limitations in a single resource.

### Remark 1

A slightly different mathematical model of translation (8) would have the mRNA/ribosome complex dissociating directly upon protein production, and would be as follows: 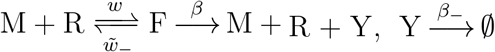. Nevertheless, the basic competition phenotype remains the same as can be seen by defining 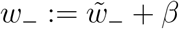 in (8).

### Remark 2

An alternative way to regulate a target circuit is to use a small molecule such as AHL to activate the target promoter [6]. The RNAP binding reactions can be written as follows (compare to (7)): 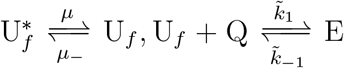, where 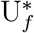 is the free inactive promoter, and *μ* is proportional to the exogenous input. our model encompasses this case also since it can be shown that the reparameterization 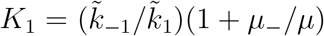 recovers the same equations (13)-(16). In other words, the effective effect of the external input in our model is to regulate the RNAP dissociation ratio *K*_1_.

### A. Limited ribosomes and abundant RNAP

In this case, RNAP Q is unaffected by the circuit (7)-(8),(9)-(10), hence its level *Q* will be constant, *Q* = *Q_T_*.

Therefore, we write the conservation law (4) as

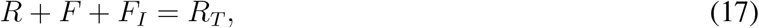

where 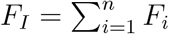.

Solving the resulting algebraic equations, the free resource at steady state is found to be:

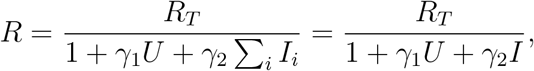

where *γ_j_* := (*A_j_Q_T_*)/(*W_j_*(*K_j_* + *Q_T_*)), *j* = 1, 2. The output is

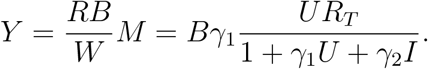

The output is linearly dependent on *R_T_*, and takes an inhibiting Michaelis-Menten form with respect to the competition *I*. It also takes a Michaelis-Menten form with respect to *U*. The CPDR *ρ* is given as: *ρ* = (1 + *γ*_1_*U*)/(1 + *γ*_1_*U* + *γ*_2_*I*), which is *independent* of the total resource *R_T_*.

We next study the properties of the output as it depends on the input and the competition.

As a function of the resource *R_T_*, the output is linear as is noted above, while it is concave with respect to *U*. As a function of the competition, it is convex. Note *d*^2^*Y/dI*^2^ > 0 for all *I* > 0.

We plot a few numerical simulations with *A_j_* = 10, *B* = 10, *W_j_* = 0.1, *R_T_* = 1, *K_j_* = 0.1, *U* = 1, *j* = 1, 2 in Figure 2 to depict the typical behavior of the competition phenotype associated with this mode of competition.

**Fig. 2.**
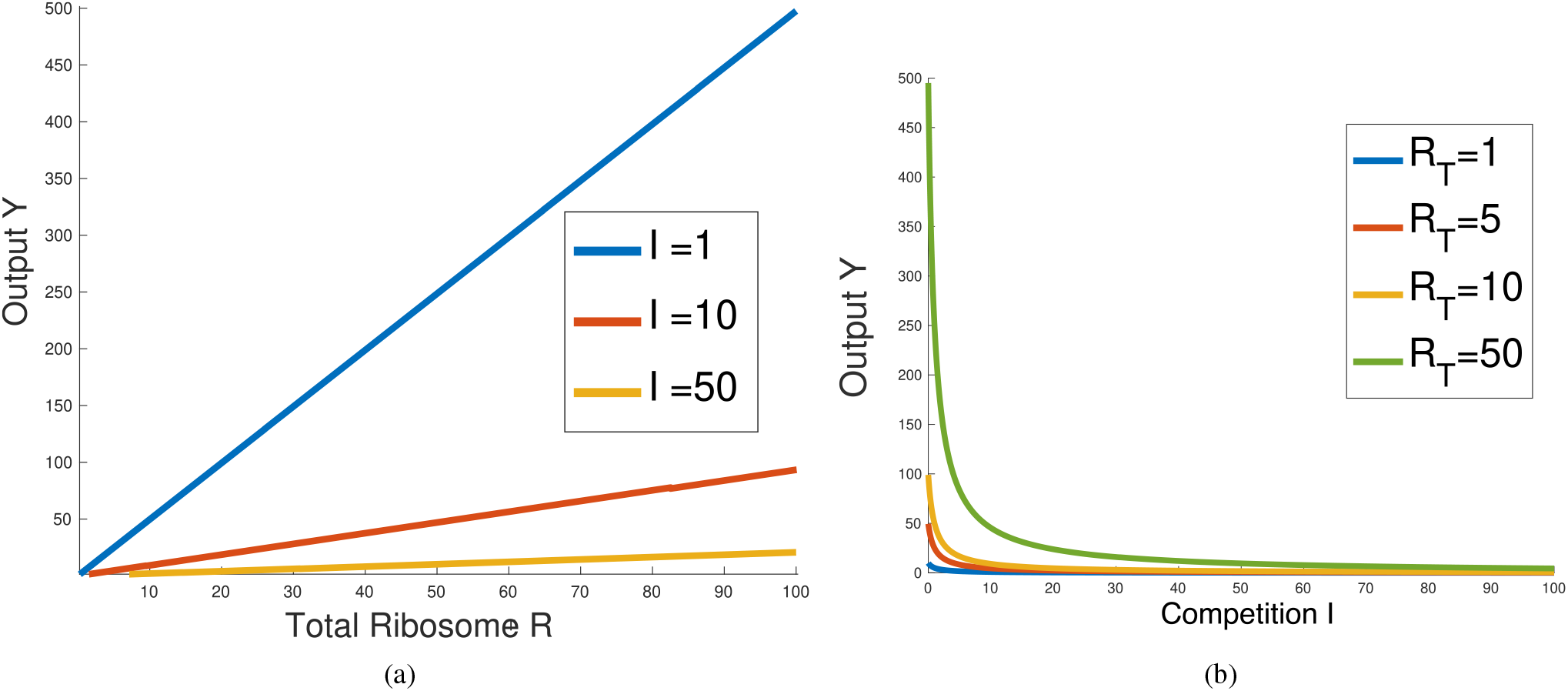
The limited ribosome case. (a) The output versus total ribosome for various competition levels, (b) The output versus total competition for various total ribosome levels.

#### Remark 3

The fact that the CPDR is independent of the total resource might lead one to think that the total resource is irrelevant for reducing competition. However, this depends on how we define competition reduction. If we consider strategies to reinstate the level of the output *Y* to its competition-free level *Y* |_*I*=0_, then we can increase the total resource to compensate for the reduction in the output due to competition. In particular, we can write the required total resource as 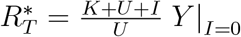.

### B. Limited RNAPs and abundant ribosomes

In this case, the ribosome R will be unaffected by the circuit (7)-(8),(9)-(10), hence its level *R* will be assumed to be constant. In other words, we have *R* = *R_T_*. Therefore, we write the conservation law (4) as:

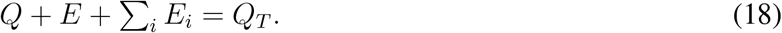

In the most general case, solving (18) for *Q* requires solving a cubic equation. Therefore, in this subsection, we assume that *K*_1_ = *K*_2_ = *K* to simplify the analysis. In this case, the free RNAP *Q* and the output *Y* are

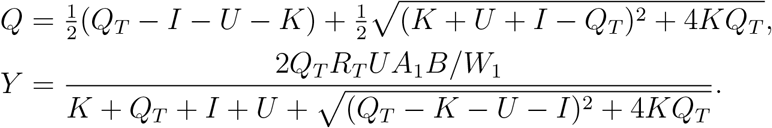

As *Q_T_* grows without bound, we have 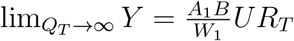, i.e., the protein is expressed at maximum capacity. If *I* grows without bound, then lim_*I*→∞_ *Y* = 0. The competition reduction ratio is written as follows:

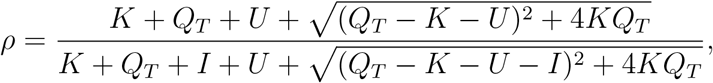

which depends on the total resource unlike the case in §2.1.

#### Convexity of the output as a function of the resource

It is globally concave as revealed by the following calculation:

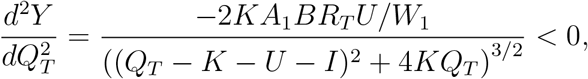

for all *Q_T_* ≥ 0. We plot few numerical simulations with *A*_1_ = 10, *B* = 10, *W*_1_ = 0.1, *R_T_* = 1, *K* = 0.1, *U* = 1 in Figure 3-a,b.

**Fig. 3.**
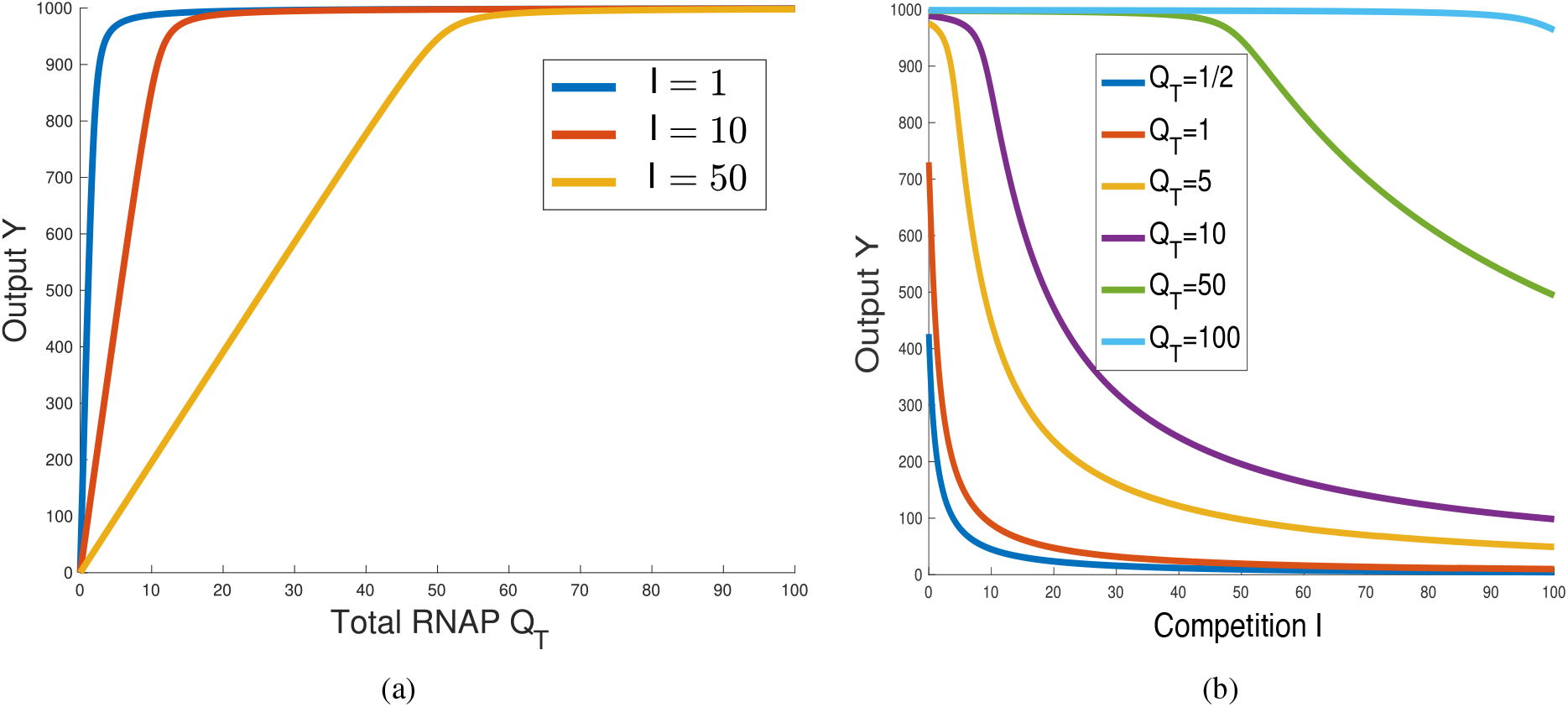
Limited RNAP case. (a) The output versus total RNAP for various competition levels, (b) The output versus competition with various *Q_T_*.

#### Convexity of the output as a function of the competition

Simulations show that the response is globally convex when *Q_T_* is small. For larger *Q_T_*, the response starts concave and then it has an inflection point. Using the same parameters above, when *Q_T_* = 1/2 the response is convex at zero as verified by computing the second derivative of *Y* with respect to *I* at zero. Figure 3-c shows the transition from convexity to concavity with higher *Q_T_*.

### C. Distinguishing the competition phenotypes

#### a) Convexity/concavity

We have been able to prove that, at steady state, limited ribosome and limited RNAP result in *qualitatively distinct competition phenotypes*. For the second the output takes a Michaelis-Menten form with respect to the resource (RNAP) level, but, in contrast, for the first it is perfectly linear with respect to the resource (ribosome level); see Fig. 2. Furthermore, a high level of RNAP provides *buffering* against low levels of competition in the second case, while the output drops quickly even with low competition in the first case; see Fig. 2b. We are able to characterize this mathematically by proving that the output is initially concave (i.e., superlinear) with respect to competition mediated by limited RNAP, while it is globally convex (i.e., sublinear) with limited ribosomes. Thus, using *either* criterion (titrating resource, or titrating competitors), we can see a clear difference between these two types of context limitations. From a different point of view, theory helps us *guess the source of competition* based upon experimental data.

#### b) Effect of the RNAP dissociation ratio

The effective RNAP DR *K*_1_ can be modified by using an inducible promoter (see Remark 2). Keeping the remaining parameters fixed, we may think of the outputs *Y* and 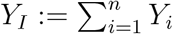 as functions of *K*_1_. Let us analyze now the trade-off between the two outputs when varying the parameter *K*1 while keeping the resource levels constant. The resulting parametric curves are also known as *isocost curves* in the language of economics [6].

In the case of limited ribosomes and abundant RNAP (i.e., *Q* = *Q_T_*), it can be shown that the relationship between the parameterized outputs is *linear* and is given by the following parametric equation:

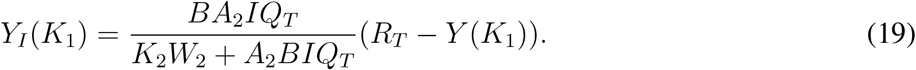

Note that (19) is *independent* of the ribosome DR *W*_1_.

In comparison, the case of limited RNAP and abundant ribosomes (i.e., *R* = *R_T_*) is more complicated. The corresponding relationship between *Y* (*K*_1_) and *Y*(*K*_1_) is generally *nonlinear* and its computation requires solving a cubic equation, as noted before. In order to probe the effect of *K*_1_, we use the simplifying assumption *Q* ≪ *K*_1_, *K*_2_ which is often satisfied by practical systems [6]. Under this approximation, it can be shown again that the relationship between *Y*(*K*_1_) and *Y_I_* (*K*_1_) is linear and is given by the parametric equation:

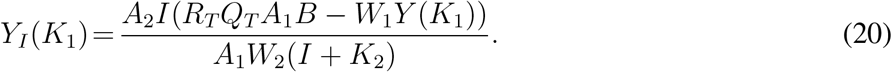

In this case, the relationship depends on *W*_1_. Therefore, the two modes of competition (for RNAP or ribosomes) are in principle distinguishable by modifying *W*_1_ via variable Ribosome Binding Site (RBS) strengths.

Even though the linear approximation (20) may not hold in all situations, dependence on *W*_1_ in the *K*_1_-parameterized relationship between *Y* and *Y_I_* indicates limited RNAP as seen in Figure 4-a) which depicts a simulation of the parameterized curves for various values of *W*_1_. In comparison, Eq. (19) is linear and is independent of *W*_1_. Figure 4-b) shows experimental data from [6] that depicts the same scenario. Our theoretical and computational prediction is consistent with the slope change noticed in the experimental data. We derived our conclusions assuming that RNAP is limited and that ribosomes are abundant. In practical situations, both resources might be in limited supplies [6]..

**Fig. 4.**
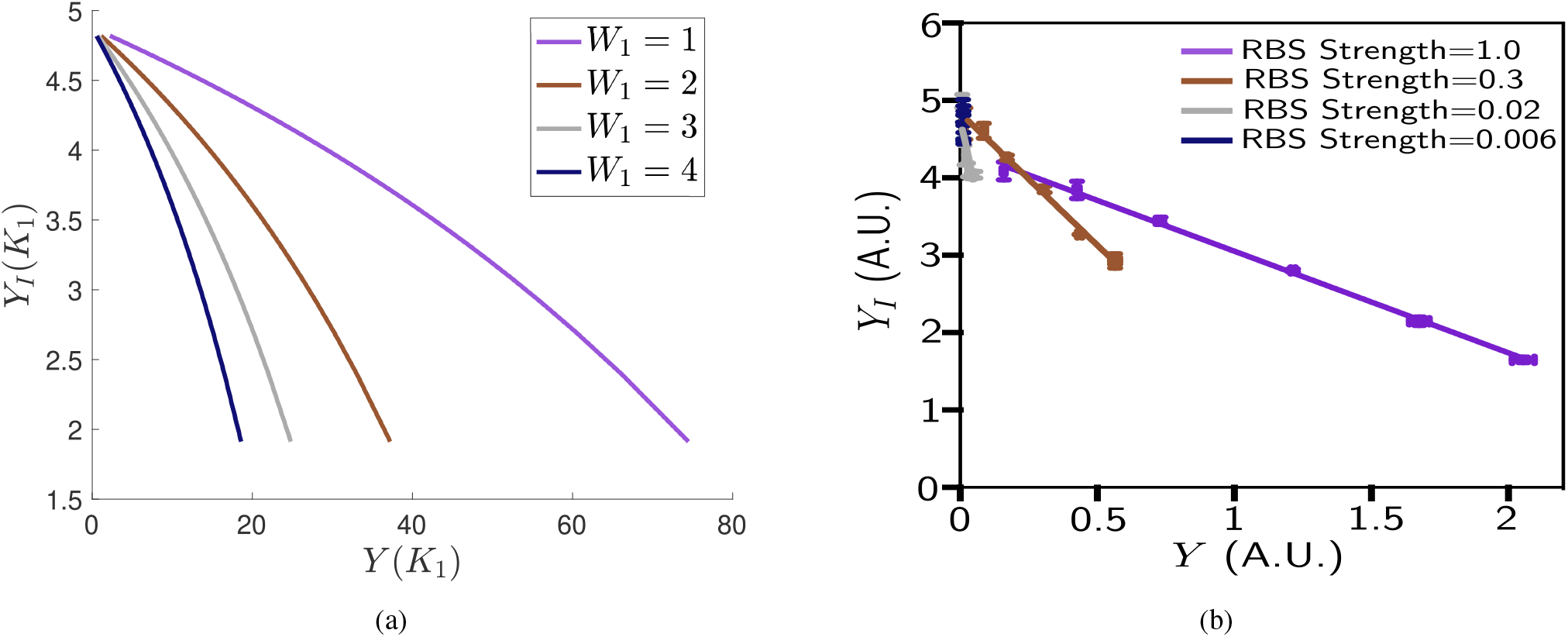
Limited RNAP manifests as sensitivity to the ribosome DR *W*_1_ when parameterizing the outputs in terms of the RNAP DR *K*_1_. (a) Simulated (*Y*(*K*_1_), *Y_I_*(*K*_1_)) parametric curves. The curves are computed by solving the cubic equation (18). The parameters are: *U* = 10,1 = 10, *W*_1_ = 4, *A*_1_ = *A*_2_ = 1, *B* = 1, *K*_2_ = 10, *W*_2_ = 1, *Q_T_* = *R_T_* = 10, *K*_1_ ∈ [0,400]. (b) Data adapted from [6] where the curves are parameterized via the AHL concentration which corresponds to *K*_1_ in our model (see Remark 2), and by varying the RBS strengths which are inversely proportional to the phenomenological parameter *W*_1_ in our model.

#### c) Effect of the Ribosome Dissociation Ratio

Let us now consider modifying *W*_1_, instead of *K*_1_. We can again write the outputs as *Y*(*W*_1_), *Y_I_* (*W*_1_). In the case of limited Ribosomes and abundant RNAP, the relationship is again linear and it can be written in the same form as (19). It can be seen that it is independent of the RNAP DR *K*_1_.

In the case of limited RNAP and abundant ribosomes, the free RNAP *Q* which solves Eq. (18) is independent of ribosome DR *W*_1_. Hence, the expressions *Y* (*W*_1_), *Y_I_* (*W*_1_) are not related, and *Y_I_* (*W*_1_) is a *constant.* Under the assumption *Q* ≪ *K*_1_, *K*_2_, we get:

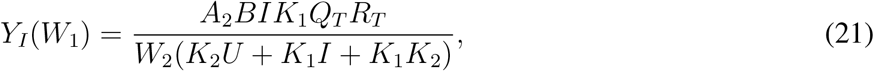

which depends on the RNAP DR *K*_1_.

Figure 5-a) shows that the case of limited ribosomes manifests as a decreasing linear relationship when parameterized by *W*_1_ (which can be experimentally controlled by RBS strengths), but it is independent of *K*_1_ (which can be controlled by utilizing an inducible promoter). On the other hand, the same relationship is constant in the case of limited RNAP, but the level is dependent on *K*_1_. This manifests by examining the *Y_I_*-axis intercepts which depend on *K*_1_ in the case of limited RNAP and abundant ribosomes, but are independent of it in the case of limited Ribosomes and abundant RNAP.

**Fig. 5.**
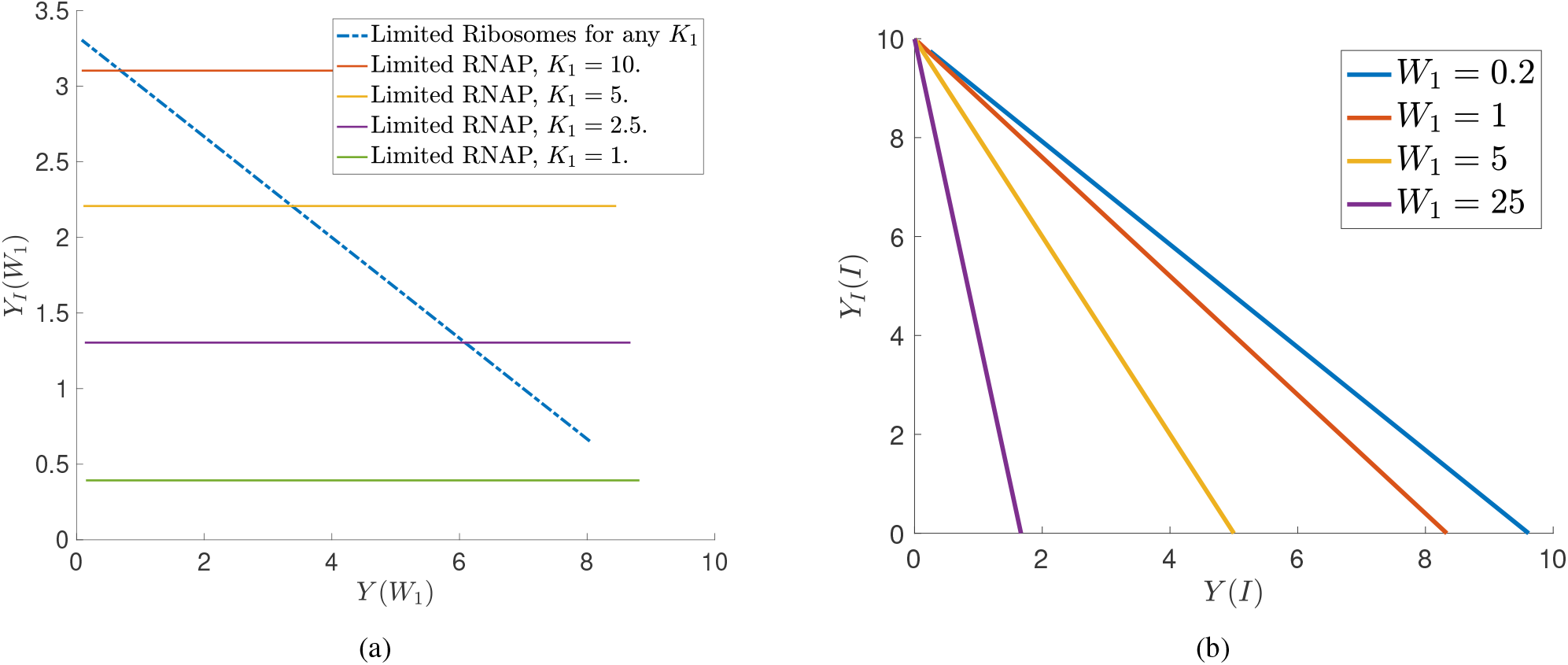
Various interventions can be used to distinguish between limited RNAP versus limited Ribosome cases. (a) simulated (*Y*(*W*_1_), *Y_I_*(*W*_1_)) curves for various *K*_1_ levels. The parameters are: *U* = 10, *I* = 1, *W*_1_ = 4, *A*_1_ = *A*_2_ = 1, *B* = 1, *K*_2_ = 10, *W*_2_ = 1, *Q_T_* = *R_T_* = 10, *W*_1_ ∈ [0,400]. (b) Simulated (*Y*(*I*), *Y_I_*(*I*)) curves for various *W*_1_ levels for the case of limited ribosomes and abundant RNAP. The parameters are similar to panel (a) with *K*_1_ = 10.

#### d) Effect of the total copy number of the competitor

We consider next modifying *I* while keeping *U* fixed. We write the outputs as *Y*(*I*),*Y_I_*(*I*).

In the case of limited ribosomes and abundant RNAP, we get a *linear* relationship as follows:

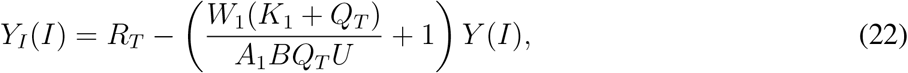

where the *Y_I_*-axis intercept depends only on *R_T_*.

In the case of limited RNAP and abundant ribosomes, the relationship is generally nonlinear, but under the assumption *Q* ≪ *K*_1_,*K*_2_ we get the following linear relationship:

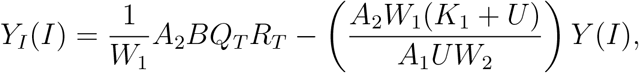

where the *Y_I_*-axis intercept depends on the ribosome DR *W*_1_. Figure 5-b) depicts the parametric curves corresponding to the limited ribosome curve where it can be seen that the *Y_I_*-axis intercept is constant for various *W*_1_.

#### e) Summary

The results are summarized in Table I.

**TABLE I.** Comparison of the competition phenotypes discussed in §2.

|  | **Limited Ribosomes**<br>**and Abundant RNAP** | **Limited RNAP and**<br>**Abundant Ribosomes** |
| --- | --- | --- |
| Resource<br>Convexity | globally linear | globally concave |
| Competition<br>Convexity | globally convex | globally convex<br>(low resource)<br>concave then convex<br>(high resource) |
| $Y_I$ vs $Y$<br>parameterized by<br>RNAP DR ($K_1$) | Linear.<br>Independent of $W_1$ | Nonlinear. Depends on<br>$W_1$ |
| $Y_I$ vs $Y$<br>parameterized by<br>Ribosome DR ($W_1$) | Linear.<br>Independent of $K_1$ | Constant. Depends on<br>$K_1$ |
| $Y_I$ vs $Y$<br>parameterized by<br>competitor copy<br>number ($I$) | Linear.<br>Slop depends on<br>$W_1$ | Nonlinear.<br>Both slop & intercept<br>depend on $W_1$ |

## III. Transcription/Translation model: Ribosome does not protect mRNA

In many situations, ribosomes do not protect mRNAs completely from decay [14]. To model this, we modify (7)-(8),(9)-(10) by adding the following reactions which describe the decay of mRNA while bound to the ribosome:

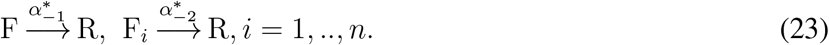

The network is no longer detailed balanced after the inclusion of decay of the bound mRNA-ribosome complexes. The steady-values of *M, M_i_* are now dependent on the resource *R*. But we still have the promoter-RNAP complex expressed at steady-state as:

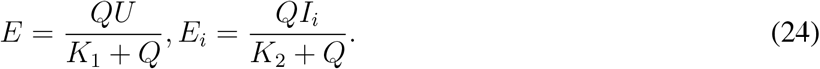

Let 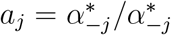 be the ratio of the decay rates of the bound and unbound mRNA. The steady-state values of the mRNAs are given as:

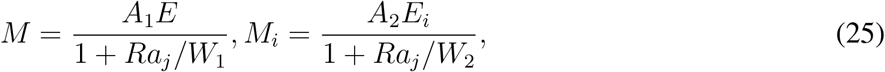

where 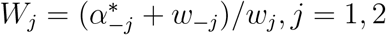.

The steady-values of the mRNA-ribosome complexes and the output in terms of the free resource *R* are given as:

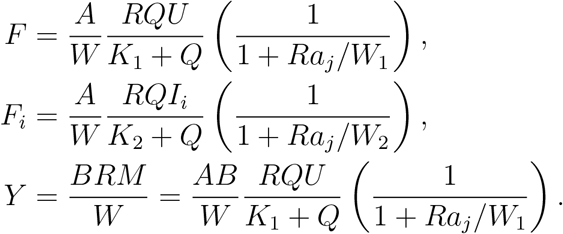

Observe that when 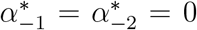, we recover the case discussed in §2. We study different scenarios next.

### A. Limited ribosomes and abundant RNAP

As before, we have abundant RNAP, hence *Q* = *Q_T_*. The only conservation law is (17).

Therefore, we need to solve for the free ribosome *R*. Let *E* be as given in (24), and let *E_I_* = ∑_*i*_ *E_i_* where *E_i_* is defined in (24). The conservation law (17) results in a cubic equation. Therefore, we assume that *W* = *W*_1_ = *W*_2_, *K* = *K*_1_, *K*_2_ in order to simplify the analysis. We get

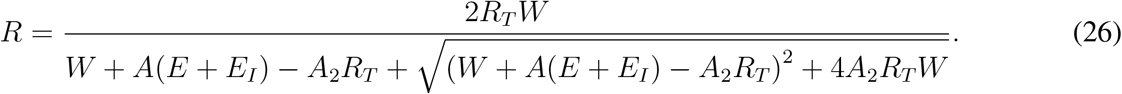

The output can be written as in:

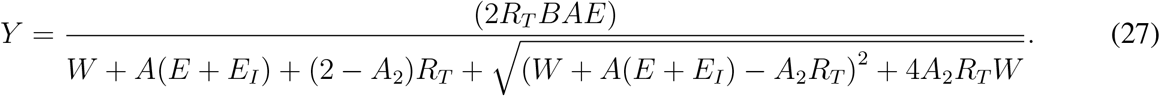

Note that when *a*_1_ = *a*_2_ = 0, we get the case discussed in §2.1. Next, we show that the properties of the system above can be deduced by studying a different system that has been studied earlier.

### B. Different systems have the same competition phenotype

It is perhaps surprising, that *two very different biological systems may lead to mathematically identical competition phenotypes.* To formalize this, let us write the (6) as: *Y* = *H*(*V*; Λ) where the inputs are *V* = [*U, R_T_,I*_1_,..,*I_n_*]^*T*^, and Λ are a set of parameters (kinetic rates, for example) that appear in (1)-(3). Hence, we have a steady-state expression *H*_1_(*V*_1_;Λ_1_) that gives us the amount of output, as well as a second function *H*_2_(*V*_2_;Λ_2_) for a different system; equivalent phenotypes will have the property that there is a diffeomorphism *ψ*: (*V*_1_, Λ_1_) ↦ (*V*_2_, Λ_2_) so that every function *H*_2_(*V*_2_;Λ_2_) can be written as *H*_1_(ψ(*V*_1_, Λ_1_)).

As a concrete example, let us revisit scenarios §2.2 and §3.1 discussed earlier with *A*_1_ = *A*_2_ = *A*. For the system in §3.1, we consider the case in which the mRNA decays at the same rate, whether it is bound to the ribosome or not, i.e., *a*_1_ = *a*_2_ = 1. One can prove that the two systems are equivalent, under a reparameterization given as follows:

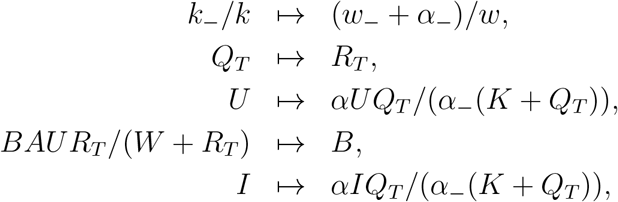

where *I* := ∑_*i*_ *I_i_*. Note that the underlying biochemical systems are very different, and the two systems of oDEs are distinct. In fact, they result in different transient behavior. However, the steady-states are the same, as shown theoretically and illustrated in Fig. 6.

**Fig. 6.**
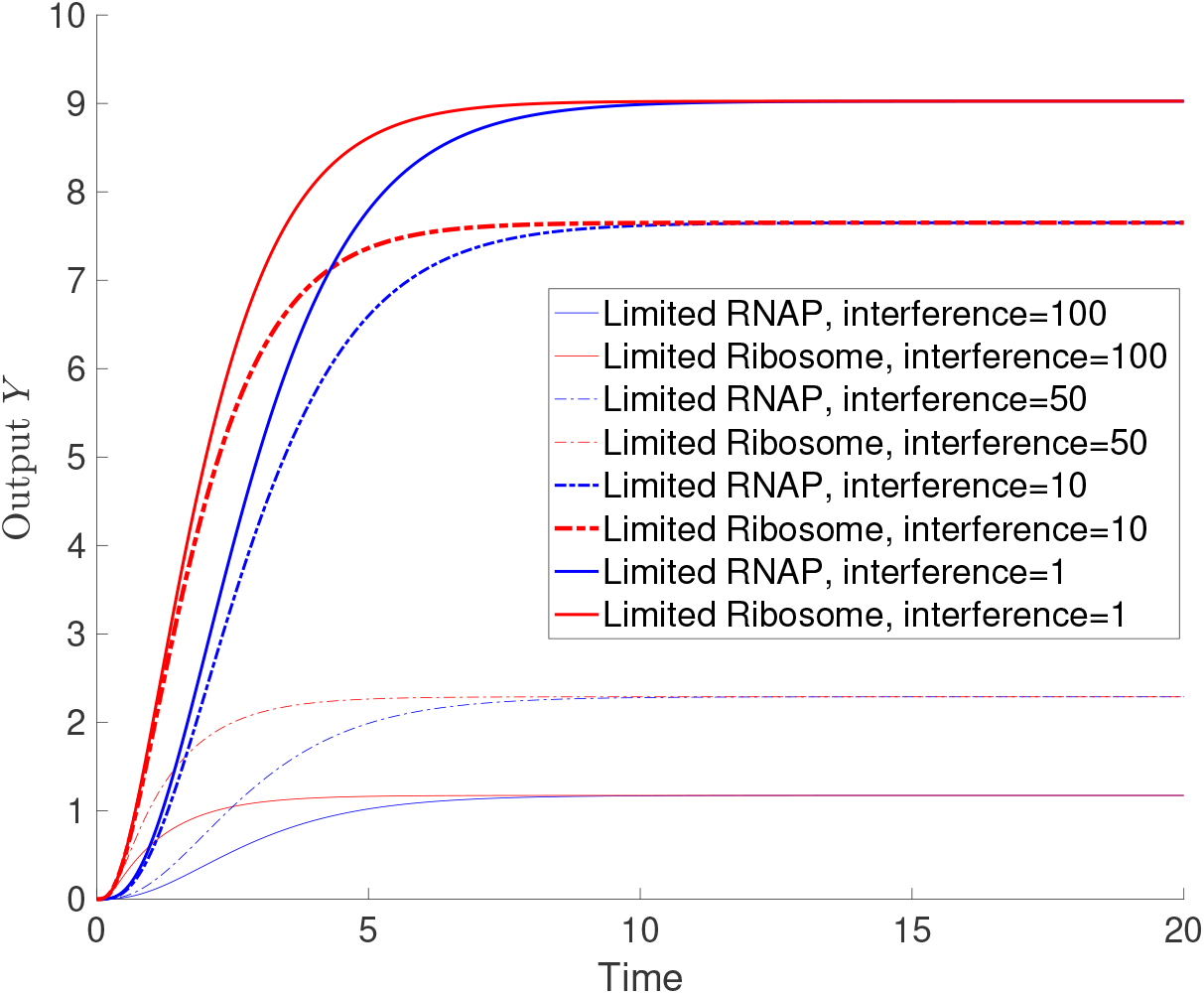
Two different competition systems with different transient are identical at steady state.

### C. Abundant ribosome and limited RNAP

Similar to the previous section, we let *R* = *R_T_*. Hence, the only conservation law is (18). Since decay affects only the ribosome-mRNA complex, we can see immediately that this case is very similar to the case discussed in §2.2 except for an additional factor. Hence, similar analysis can be replicated.

## IV. Beyond Transcription/Translation systems

The competition phenotypes studied in the previous sections can be generalized to other biological contexts by classifying them into two main categories:

### A. Externally-regulated targets

The case discussed in §2.1 has the ribosome as the limited resource, hence the target that consumes the resource is mRNA which is not conserved. Hence, the model can be written essentially as follows:

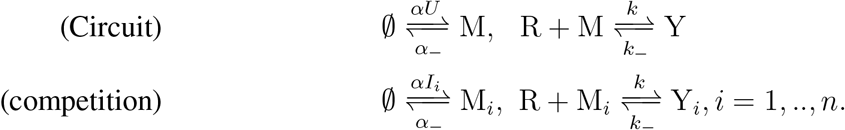

In addition to mRNAs, the above model can represent ligand-activated enzymes M, M_*i*_, *i* = 1,.., *n* competing for a substrate R, single-guide RNAs (sgRNAs) competing for a limited amount of dCas9 in CRISPRi [15, 16], or externally-activated TFs competing for a single promoter.

### B. Conserved targets

The case discussed in §2.2 has the RNAP as the limited resource, hence the target that consumes the resource is the promoter which is conserved. Hence, the model can be written essentially as follows:

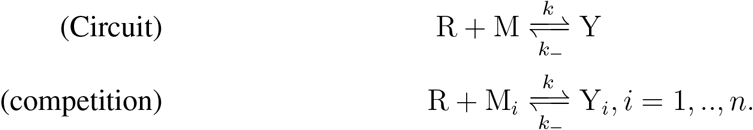

In addition to promoter, the above model can represent conserved enzymes M, M_*i*_, *i* = 1,..,*n* competing for a substrate R, or conserved TFs competing for a single promoter.

